# The complete plastid genomes of four species from Brassicales

**DOI:** 10.1101/458000

**Authors:** Weixue Mu, Ting Yang, Xin Liu

## Abstract

Brassicales is a diverse angiosperm order with about 4,700 recognized species. Here, we assembled and described the complete plastid genomes from four species of Brassicales: *Capparis urophylla* F.Chun (Capparaceae), *Carica papaya* L. (Caricaceae), *Cleome rutidosperma* DC. (Cleomaceae), and *Moringa oleifera* Lam. (Moringaceae), including two plastid genomes newly assembled for two families (Capparaceae and Moringaceae). The four plastid genomes are 159,680 base pairs on average in length and encode 78 protein-coding genes. The genomes each contains a typical structure of a Large Single-Copy (LSC) region and a Small Single-Copy (SSC) region separated by two Inverted Repeat (IR) regions. We performed the maximum-likelihood (ML) phylogenetic analysis using three different data sets of 66 protein-coding genes (ntAll, ntNo3rd and AA). Our phylogenetic results from different dataset are congruent, and are consistent with previous phylogenetic studies of Brassiales.

---

Brassicales, an order of flowering plants that contains ~4,700 species (Kiefer et al. 2014; Cardinal-McTeague et al. 2016), is a monophyletic group consisting 17 families including several model families, such as Brassicaceae, for development and evolutionary biological studies (Soltis et al. 2011; Chase et al. 2016). Brassicales are known for their rich diversities in morphology, physiology, development, and chemical traitse (Koenig and Weigel, 2015). Here, we reported the complete plastid genomes of four Brassicales species that were assembled using the Next-Generation Sequencing approach with BGISEQ-500 (BGI, Shenzhen) paired-end reads.

In this study, the samples were collected from Ruili Botanical Garden at Yunnan, China. The voucher specimens have been deposited in the Herbarium of China National GeneBank (HCNGB). The fresh leaves of each sample were used to extract total genomic DNA using the modified CTAB method (Doyle and Doyle 1987). High-throughput sequencing was carried out using an BGISEQ-500 platform. Approximately 50 Gb high quality, 100 base pairs (bp) paired-end reads were obtained for each sample. The raw reads with low quality were filtered out using the program SOAPfilter_v2.2. *De novo* assemblies were carried out using the seed-extension-based *de novo* assembler NOVOPlasty v2.5.9 (Dierckxsens et al. 2016). A complete rbcL gene sequence of *Arabidopsis thaliana* (downloaded from NCBI, accession number U91966) was used as the seed to conduct the assemblies. The longest contig of each sample were further blasted against the chloroplast database (downloaded from NCBI, including 2,503 non-redundant species), and the resulted best-hit sequences (minimum requirement: e-value < 10-7 and identity>95%) were used as the reference for further assembly using an ‘baiting and iterative mapping’ approach assembler MITObim v1.8 (Hahn et al. 2013). Complete plastid genomes were recovered for all the four species. Plastid genomes were then annotated on the standard web-based program GeSeq (Tillich et al. 2017), and the annotation of protein-coding genes were further conducted with GeneWise v2.4.1 (Birney et al. 2000). The complete plastid genomes have been submitted to the CNGB Nucleotide Sequence Archive (CNSA : https://db.cngb.org/cnsa; accession number: CNA0000813 (*Capparis urophylla* F.Chun); CNA0000814 (*Carica papaya* L.); CNA0000815 (*Cleome rutidosperma* DC.); CNA0000816 (*Moringa oleifera* Lam.).)

The assembled four Brassicales plastid genomes ranged from 155,359 bp (*Capparis urophylla* F.Chun) to 163,131 bp (*Moringa oleifera* Lam.) in length, and all share a typical quadripartite structure: two inverted repeats (IRs), a large single-copy region (LSC), and a small single-copy region (SSC). As for *Carica papaya* L. and *Cleome rutidosperma* DC., the total length of the plastid genomes are 161,752 bp and 158,477 bp, respectively. The plastid genome of *Carica papaya* L. has the longest LSC region of 130,436 bp, along with two 3,483 bp IR regions and a shortest 24,350 bp SSC region, while the plastid genome of *Capparis urophylla* F. Chun has the shortest LSC (84,518 bp), two IRs (10,318 bp) and a longest SSC (50,205 bp), respectively. The length of LSC, IR and SSC regions of *Cleome rutidosperma* DC. and *Moringa oleifera* Lam. are 107,763 bp, 2,888 bp, 45,028 bp and 102,342 bp, 3,710 bp, 48,715 bp, respectively. All four species have the same number of 78 protein-coding genes, four ribosomal RNAs, while the number of transfer RNAs varies from 36 (*Cleome rutidosperma* DC.), 37 (*Carica papaya* L. and *Moringa oleifera* Lam.) to 38 (*Capparis urophylla* F.Chun).

We performed phylogenetic analyses with all four newly assembled plastid genome sequences together with 16 published plastid genomes of Brassicales species and two Malvales species. The plastid sequences were aligned with MAFFT V7.017 (Katoh et al. 2013) on the base following three data sets: (1) all nucleotide positions (ntAll); (2) the first and second codon positions (ntNo3rd); (3) the amino acid (AA) of 78 protein-coding genes (Ruhfel et al. 2014). The Maximum-Likelihood trees were constructed under the GTRCAT (PROTGAMMAWAG for AA) model using the parallel version of RAxML v8.2.4 (Stamatakis 2014) with 100 bootstraps. The phylogenetic results revealed that the relationships constructed using the three different data sets were highly consistent at the family level (Figure 1). The plastid phylogeny recovered in our study is congruent with the previous phylogenetic study of Brassiales (APG 2016).

**Figure 1.**
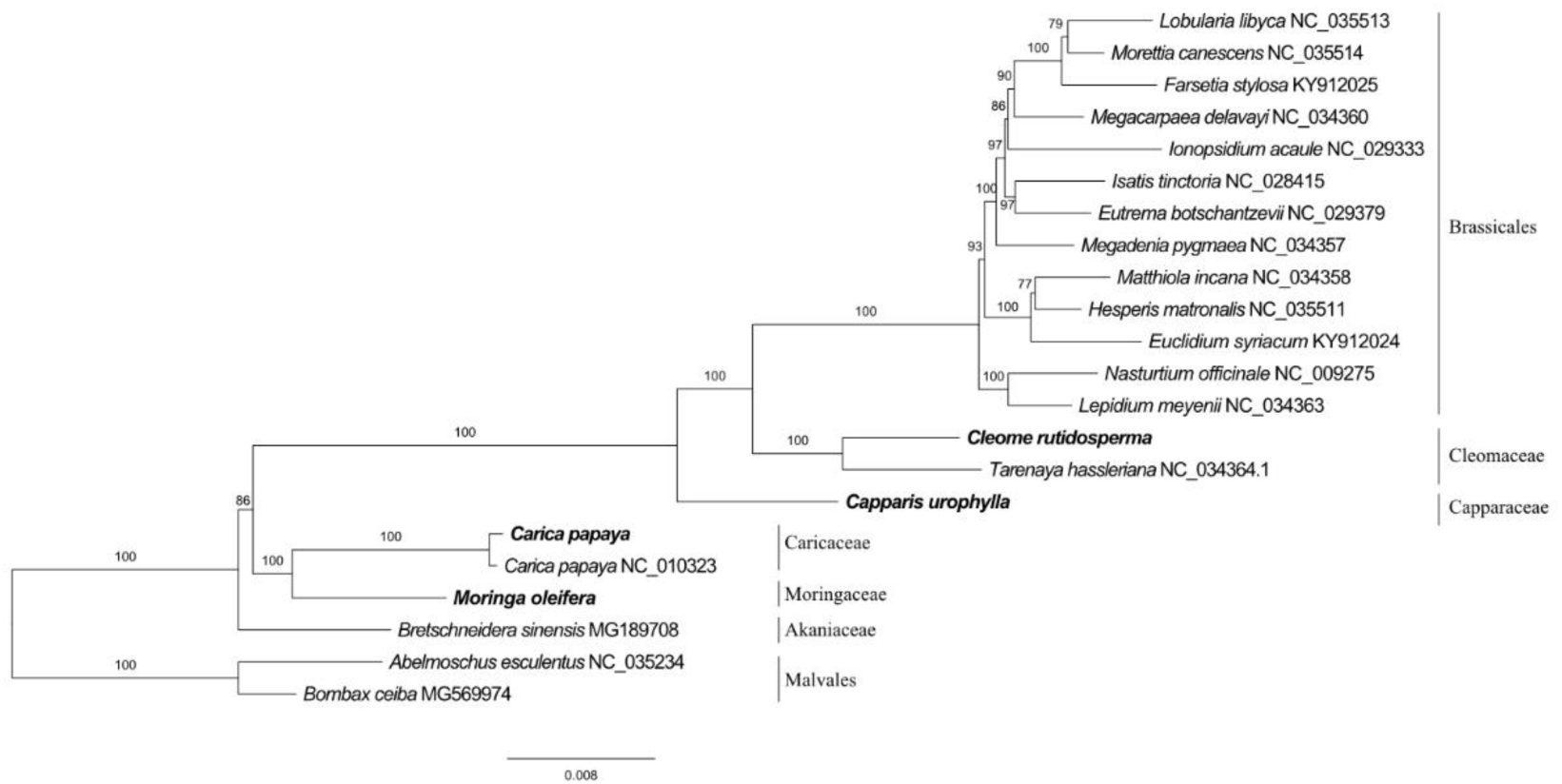
Maximum-likelihood phylogenetic tree based on 22 ntAll data sets. The numbers on the nodes indicate bootstrap support values.

